# Asiatic acid improves mitochondrial function, activates antioxidant response in the mouse brain and improves cognitive function in beta-amyloid overexpressing mice

**DOI:** 10.1101/2024.02.21.581270

**Authors:** Samantha Varada, Steve R Chamberlin, Lillie Bui, Mikah S Brandes, Noah Gladen-Kolarsky, Christopher J Harris, Wyatt Hack, Barbara H Brumbach, Joseph F Quinn, Nora E Gray

## Abstract

Extracts of the plant *Centella asiatica* can enhance mitochondrial function, promote antioxidant activity and improve cognitive deficits. Asiatic acid (AA) is one of the constituent triterpene compounds present in the plant. In this study we explore the effects of increasing concentrations of AA on brain mitochondrial function, antioxidant response and cognition in healthy mice and a single concentration of AA in the beta-amyloid overexpressing 5xFAD mouse line. Associative memory and overall activity were assessed. Hippocampal mitochondrial bioenergetics and the expression of mitochondrial and antioxidant response genes was determined. In the 5xFAD line, total beta-amyloid plaque burden after AA treatment was also evaluated. In healthy mice, we report dose responsive effects of increasing concentrations of AA on enhanced associative memory and a dose dependent increase in basal and maximal mitochondrial respiration, mitochondrial gene expression and antioxidant gene expression. Results from the highest AA dose (1% AA) were similar to what was observed with CAW. The high AA dose was then evaluated in the context of Aβ accumulation in 5xFAD mice. Improvements in mitochondrial and antioxidant response genes were favored in females over males without significant alleviation of Aβ plaque burden.

## Introduction

Alzheimer’s disease (AD) is the most common neurodegenerative disease of the elderly currently estimated to affect 6.7 million people in the United States and predicted to affect more than twice that number by 2050 [1]. The pathophysiology of AD is characterized by extracellular amyloid-β (Aβ) plaque deposits and intracellular neurofibrillary tangles leading to neuronal loss. These changes are associated with the clinical symptoms of memory loss, cognitive decline and inability to perform daily living activities [2, 3]. AD is terminal and the development of novel therapies is hindered in part by an incomplete understanding of the mechanisms that drive disease onset and progression.

Mitochondrial dysfunction and oxidative stress (OS) are early events in the AD brain that are believed to contribute to the progression of the disease [4]. The magnitude of metabolic activity in the brain. coupled with limited energy reserves, create an environment with high levels of reactive oxygen species (ROS) that must be balanced by appropriate antioxidant response to avoid increased generation of OS. During aging and in AD, antioxidant response declines while at the same time mitochondrial dysfunction increases [5]. In fact in AD, OS and mitochondrial dysfunction are early events that precede the onset of other symptoms [5, 6]. The mitochondrial changes include impaired bioenergetics as well as reduction in the expression of mitochondrial enzymes in the electron transport chain (ETC) [7]. Mitochondria are both the main source of and target for ROS. Normal oxidative metabolism generates ∼90% of intracellular ROS, the levels of which are tightly regulated by antioxidant enzymes [8]. However, mitochondrial dysfunction results in even greater production of ROS which when coupled to the decreased antioxidant response observed in AD contributes to neuronal degeneration [9]. Therefore, identifying therapies that target antioxidant response and mitochondrial dysfunction could prove an effective strategy for therapeutic intervention.

The endogenous antioxidant response pathway is regulated by the transcription factor, NRF2 (nuclear factor (erythroid-derived 2)-like2; also called NFE2L2), which activates the expression of cytoprotective enzyme genes through binding of the antioxidant response element (ARE) in the promoter. NRF2 expression and its ARE gene targets are downregulated in AD [6]. Induction of NRF2 has been shown to rescue cognitive deficits in AD mouse models [10–12]. NRF2 has also been shown to regulate mitochondrial genes including glucose-6-phosphate dehydrogenase, the enzymes of the pentose phosphate pathway, malic enzyme 1, and isocitrate dehydrogenase 1 [13], further linking antioxidant response and mitochondrial function.

The medicinal plant *Centella asiatica* (L.) Urban, (Apiaceae) is an herb with an extensive history of use in Ayurvedic and Chinese traditional medicine to boost memory and enhance cognitive function [14]. There have been small clinical studies that similarly observed beneficial effects of extracts of the plant in both healthy and impaired populations with no reported adverse events [15, 16]. *Centella asiatica* is also an example of a therapeutic agent that modulates mitochondrial function and the antioxidant response. Our lab demonstrated that a water extract of *Centella asiatica* (CAW) can activate NRF2 induction both *in vitro* and *in vivo,* and in mouse models of aging and AD that activation was accompanied by improved cognitive function in CAW-treated animals [11, 12, 17–22].

The complex chemical nature of whole plant extracts makes clinical translation of standardized CAW formulation challenging. *Centella asiatica* contains four triterpene compounds: asiatic acid (AA), madecassic acid, asiaticoside and madecassoside [16]. Each of these compounds has been shown to elicit antioxidant activity and improve mitochondrial function [23–33]. Although levels of AA are relatively low in CAW the concentration of its metabolic precursor asiaticoside is quite high [34]. Studies have shown that in both humans and rodents orally administered asiaticoside gets rapidly and efficiently converted to AA *in vivo* and AA is readily absorbed and detectable in the blood stream [35–39]. AA also has an excellent safety profile having been administered to humans in multiple studies without any notable adverse events [38, 40–43] underscoring its potential use as a therapeutic agent in addition to its possible utility as a means of standardizing CAW for clinical use.

The present study aims to explore the cognitive, antioxidant and mitochondrial effects of AA. We begin by comparing the effects of a range of concentrations of AA to CAW in healthy, nine-month-old CF1 mice and then go on to evaluate the effects of AA in the context of Aβ accumulation using the 5xFAD mice.

## Methods

### CAW extract

CAW was prepared as previously described [44]. Briefly, *Centella asiatica* was obtained from Oregon’s Wild Harvest (Redmond, OR) and the water extract was prepared by refluxing 160g of the raw plant material with 2L of water for 2h. This extract was filtered and lyophilized to a powder. A representative sample of both the raw *Centella asiatica* and the CAW is retained by our lab at −20C. A full description of the chemical composition, determined by targeted liquid chromatography-high resolution tandem mass spectrometry analysis can be found in our previous publication [44].

### Mouse experimental diets

Either CAW or AA (Sigma Aldrich, St Louis, MO) was incorporated into AIN-93M diet by Dyets Inc. (Bethlehem, PA, USA). CAW was incorporated at 1% w/w to approximate the maximum 1000 mg per kg of body weight per day (mg/kg/d) dose used in our previous dose-response study [21] AA was incorporated into AIN-93M at one of three concentrations: 1) 0.05% or 5 mg AA in 1kg diet to match the concentration of AA (w/w) in the 1000 mg/kg/d CAW diet (based on our previously published liquid chromatography-high resolution tandem mass spectrometry analysis of the CAW extract [44]); 2) 0.5% or 50mg AA in 1kg diet to match the concentration of AA+AS (w/w) in the 1000mg/kg/d diet [44] or 3) 1% AA ir 1000mg AA in 1kg diet. Diets were sterilized by gamma irradiation (5.0–20.0 kGy) at Sterigenics (Oak Brook, IL, USA).

### Animals

Female CF1 mice were obtained from Charles River. Animals were kept in a climate-controlled environment with a 12h light/dark cycle and provided with water and diet ad libitum until aged to 9 months old. All procedures were conducted according to the NIH Guidelines for the Care and Use of Laboratory Animals and approved by the institutional Animal Care and Use Committee of the Portland VA Healthcare System. At 9 months of age mice were taken off the standard diet and fed AIN-93M (vehicle diet), or AIN-93M containing either 1% CAW or 0.05%, 0.5% or 1% AA (n=9-12 per treatment condition). Treatment continued for a total of four weeks. In the final week of treatment mice underwent Conditioned Fear Response testing and then were euthanized and tissue was collected (Figure 1).

**Figure 1:**
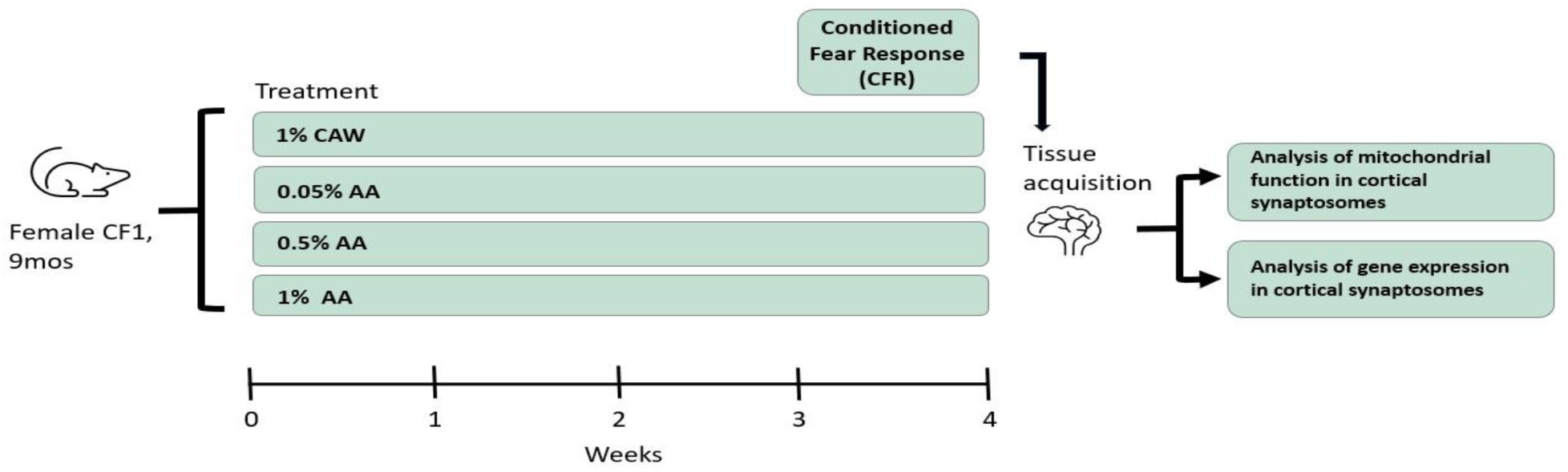
Experimental design AA dose response in CF1 mice.

5xFAD colonies were developed from breeding pairs obtained from The Jackson Laboratory. These mice overexpress human amyloid precursor protein (APP) and human presenilin 1 (PS1) with five mutations associated with familial Alzheimer’s Disease (FAD): the Swedish (K670N, M671L) Florida (I716V) and London (V717I) mutations in APP and two in PS1 (M146L and L286V) [45]. The 5xFAD line develops amyloid plaques at a young age (2 months) with cognitive impairment evident by 5-6 months [45]. Animals were kept in a climate-controlled environment with a 12h light/dark cycle and provided with water and diet ad libitum until aged to 6-7 months. All procedures were conducted in accordance with the NIH Guidelines for the Care and Use of Laboratory Animals and were approved by the institutional Animal Care and Use Committee of the Portland VA Healthcare System. At 6-7 months of age mice were taken off the standard diet and fed AIN-93M (vehicle diet), or AIN-93M containing 1% AA (n = 8-12 per treatment condition). Treatment continued for a total of four weeks. In the final week of treatment mice underwent Conditioned Fear Response testing and then were euthanized and tissue was collected (Figure 2).

**Figure 2:**
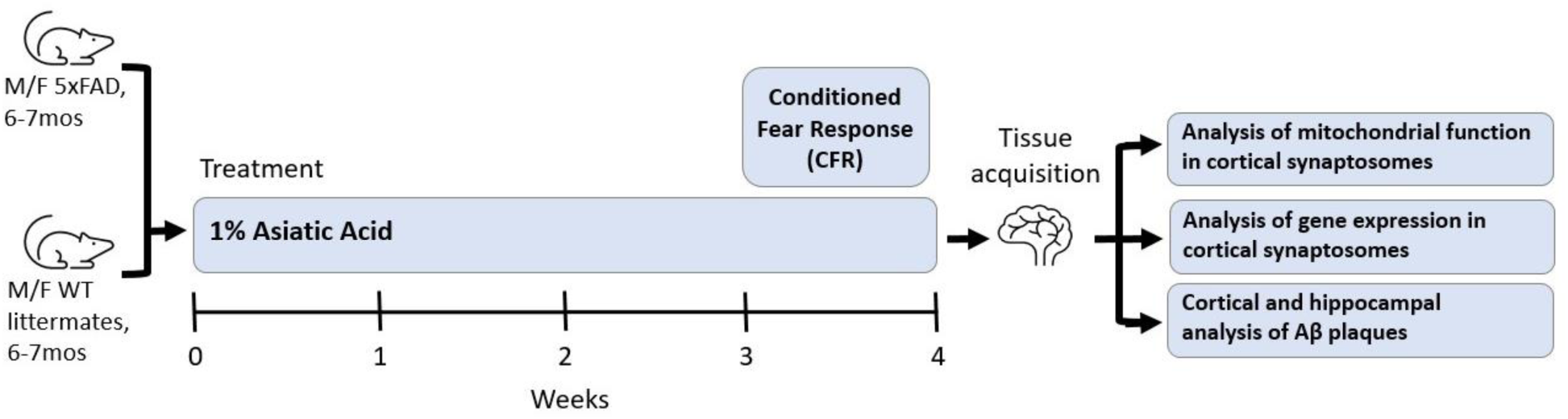
Experimental design AA treatment in 5xFAD animals.

### Conditioned Fear Response (CFR) test

The CFR test evaluates contextual memory and has been shown to be affected by inputs from the hippocampus, cortex and amygdala [46]. It has three phases: habituation, conditioning and testing. In the habituation phase an animal was exposed to a 16×16×12 inch chamber with a wire floor for 5 minutes. The conditioning phase occurred immediately following habituation, where the animal was exposed to 3 one-second shocks (0.5A) randomly distributed over a 3-minute period with no more than one shock per minute. The test phase occurred 24 hours after the conditioning phase, where the animal was reintroduced once more to the same chamber, but this time not exposed to any shocks. The amount of time spent freezing over a 10-minute period is recorded. Freezing time is represented as the change in freezing time from the habituation phase to the test phase in order to account for any baseline differences in overall activity.

### Synaptosomal isolation

Cortical tissue was harvested from the mice and synaptosomes were isolated using SynPer reagent from ThermoFisher (Waltham, MA) according to the manufacturer’s instructions. The total protein content of each synaptosomal preparation was determined by BCA assay.

### Analysis of mitochondrial function

Mitochondrial bioenergetics were quantified in isolated cortical synaptosomes taken from the left hemisphere using the Seahorse Xfe96 Analyzer. 10ug of total synaptosomal protein was plated in each well of a polyethyleneimine coated 96 well plate (n=5-6/animal) and analyzed using the MitoStress kit from Agilent (Santa Clara, CA). The MitoStress kit measures oxygen consumption rate (OCR) was measured under varying conditions. After three initial baseline measurements the ATP synthase inhibitor oligomycin (2 μM) was added and three more measurements were taken. The difference between the average of these values and the average of the basal respiration reflects respiration is the ATP-linked respiration. Next, an ETC accelerator, p-trifluoromethoxy carbonyl cyanide phenyl hydrazine (FCCP; 2 μM), was added and 3 measurements were taken. The average of these values reflects maximal respiration. The difference between maximal and basal respiration is the spare capacity. Finally the ETC inhibitors rotenone and antimycin (0.5 μM) were added.

### Gene Expression

RNA was isolated from the remainder of the synaptosomal preparations using Tri-Reagent (Thermo Fisher, Waltham, MA) using the protocol provided by the manufacturer. RNA was reverse transcribed with the Superscript III First Strand Synthesis kit (Thermo Fisher, Waltham, MA) to generate cDNA as per the manufacturer’s instructions. Relative gene expression was determined using TaqMan Gene Expression Master Mix (Invitrogen) and commercially available TaqMan primers (Invitrogen) for synaptophysin (Mm00436850_m1), post-synaptic density protein 95 (PSD95; Mm00492193_m1), NRF2 (Mm00477784_m1), glutamate-cysteine ligase catalytic subunit (GCLC; Mm00802655_m1), heme oxygenase 1 (HMOX1; Mm00516005_m1), mitochondrially encoded NADH:Ubiquinone Oxidoreductase Core Subunit 1(Mt-ND1; Hs02596873_s1), mitochondrially Encoded Cytochrome B (Mt-CYB; Hs02596867_s1), Mitochondrially Encoded Cytochrome C Oxidase I (Mt-CO1; Hs02596864_g1), Mitochondrially Encoded ATP Synthase Membrane Subunit 6 (Mt-ATP6; Hs02596862_g1) and glyceraldehyde-3phosphate dehydrogenase (GAPDH; Hs02758991_g1). Quantitative PCR (qPCR) was performed on a StepOne Plus Machine (Applied Biosystems) and analyzed using the delta-delta Ct method.

### Immunohistochemistry

In the 5xFAD experiment the right hemisphere of each animal was fixed in 4% paraformaldehyde and then passed through a sucrose gradient, and frozen. 40-micron frozen coronal sections were then cut on a freezing microtome. Sections were incubated with agitation in blocking buffer (100 mM TBS, pH 8.0, 2 mg/ml bovine serum albumin, 2% horse serum, 0.5% Triton X-100) for 2 hours, then incubated overnight with primary antibody directed against Aβ (Invitrogen, beta Amyloid (1-40) Polyclonal Antibody, # 44-136) diluted 1:1000 in blocking buffer. Sections were then incubated for 2 hours with biotinylated secondary antibody (1:200, Vector Labs, Burlingame, CA), for 2 hours with an avidin-linked peroxidase complex (ABC, Vector Labs), then developed with diaminobenzidine (DAB, Sigma) in PBS. Sections were washed, mounted in Permount (Fisher Scientific, Pittsburg, PA) and cover slipped. Protein expression was quantified in at least three coronal sections from each mouse, representing anterior, middle and posterior hippocampus and cortex. Hippocampal and cortical areas were traced using a computerized stage and stereo investigator software (Image J, Wayne Rasband, NIH, USA). Aβ plaque burden was expressed as percentage of hippocampus or cortex occupied by these plaques.

### Statistical analysis

#### AA treatment in CF1 mice

All outcome variables were assessed for normality. CFR, distance traveled, basal respiration, ATP-linked respiration and GCLC gene expression were all normally distributed. All other variables did not meet normality assumptions so a log transformation was performed and normality was established. Time immobile, maximum respiration, spare capacity, NRF2, HMOX1, Mt-ND1, Mt-CYB, Mt-CO1, and Mt-ATP6 analyses are on the log transformation.

We tested the relationship between level of AA (0%, 0.05%, 0.5%, 1%) and all outcome variables. We assessed a dose response by testing for a linear trend using the general linear model (GLM). We created an ordinal variable for AA level as the primary predictor for each outcome variable (see Figures 1-4). Additionally, we tested for specific group differences between AA levels in addition to a 1% CAW group for each outcome variable. GLM was used to test for model significance followed by post-hoc pairwise comparisons using Tukey and the least square means (lsmeans) statement in SAS. Significance was defined as p ≤ 0.05. All bar graphs show means with error bars indicating standard error of the mean with the F and p values listed corresponding to the significance of the linear trend. Analyses were performed using Excel, GraphPad Prism 6, and SAS 9.4.

#### AA treatment in 5xFAD mice

All outcome variables were assessed for normality (CFR, distance traveled, basal respiration, ATP-linked respiration, GCLC gene expression and Aβ levels, time immobile, maximum respiration, spare capacity, NRF2, HMOX1, Mt-ND1, Mt-CYB, Mt-CO1, and Mt-ATP6). Those that did not meet assumptions were log transformed. However, even with the transformation many outcome variables still violated assumptions. Therefore, we decided to use a nonparametric approach to test for group differences on all variables using the Kruskal-Wallis test. Post-hoc pairwise comparisons were assessed using Dunn’s test. All analyses assess variables in their original unit of measurement. We chose to use Kruskal-Wallis tests even for those variables that did meet assumptions for the parametric approach so that there was consistency across all outcomes for more balanced interpretation. Because the Kruskal-Wallis test uses medians to test for group differences, we use Box-and-Whisker plots to visually display the data. Analyses were performed using GraphPad Prism 6 and STATA16.

## Results

### AA treatment dose-dependently increases activity

To assess the behavioral effects of increasing concentrations of AA, we evaluated overall activity in 9-month-old CF1 mice that were administered AA integrated into their chow at either 0.05%, 0.5% or 1% diet for four weeks. Results were compared to animals given a control diet (0% AA) or a diet 1% CAW for the same amount of time. Testing occurred in the final week of treatment after which tissue was harvested. Distance traveled and time immobile were recorded as metrics of activity during the habituation phase of the CFR (when the animal is freely exploring an open area for 5 minutes).

The test for differences between groups did not reach statistical significance for either distance traveled or time immobile (Table 1). The dose response test for a linear trend in AA concentration on total distance traveled approached but did not reach statistical significance, (Figure 3A). There was however, a significant linear trend for AA level in relation to time immobile (Figure 3B) with time immobile decreasing as AA concentration increased.

**Figure 3.**
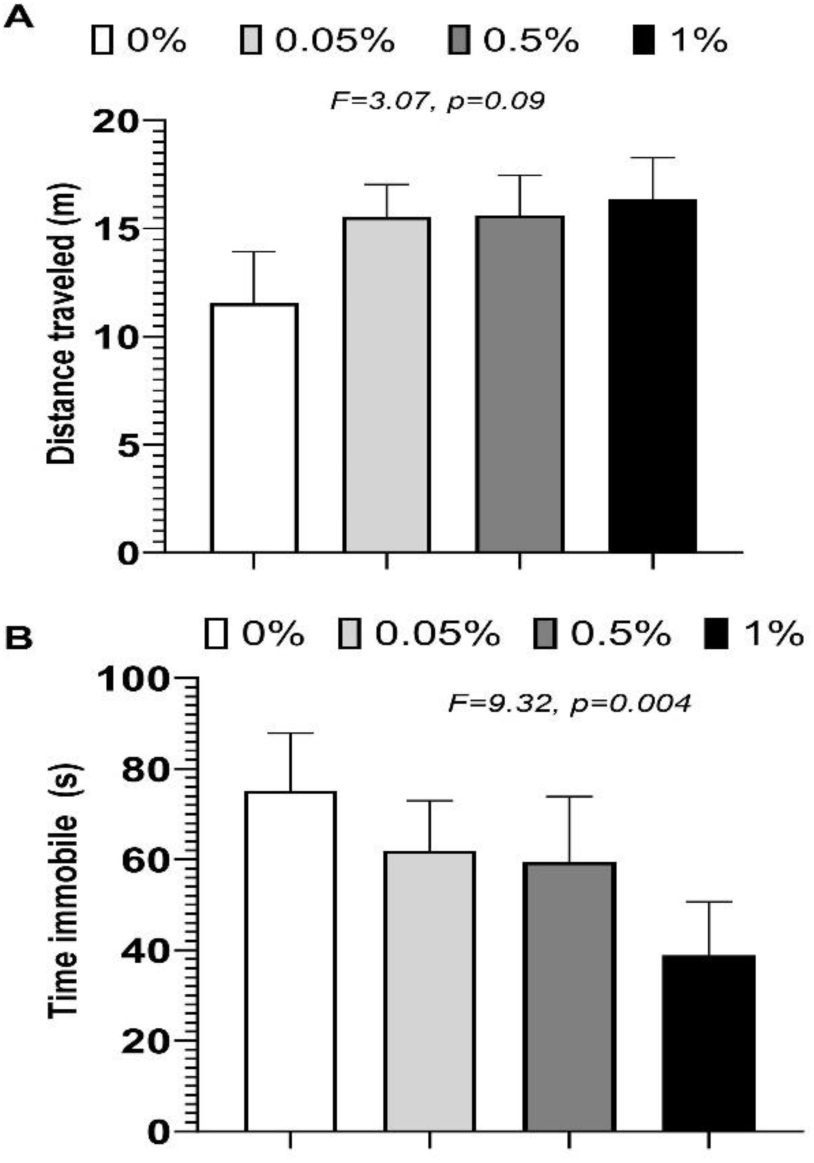
AA dose dependently increases overall mobility but not distance traveled. A) No significant linear trend was seen with increasing AA concentrations and distance traveled. B) There was a significant linear relationship between time immobile and AA concentration with higher concentrations of AA resulting in a reduction in time immobile.

**Figure 4.**
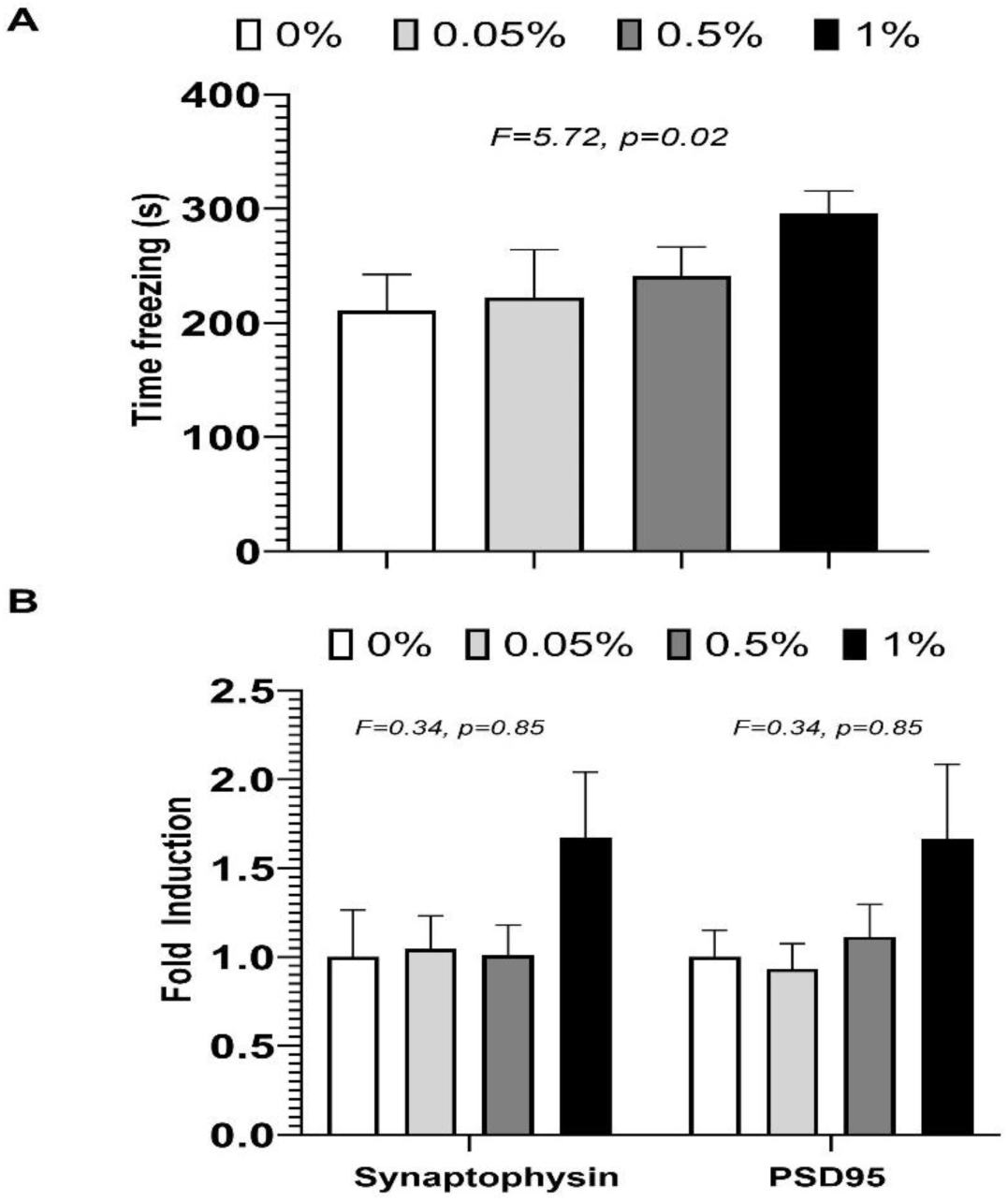
Associative memory improves in a dose-responsive manner with AA treatment. A) There was a significant linear relationship between concentrations of AA and CFR response. B) There was no significant linear relationship between expression of synaptophysin or PSD85 and concentration of AA.

**Table 1.** Group differences between AA levels and 1% CAW. Means and standard deviations are reported for all outcomes all groups. The general linear model was used to test for group differences between the control group (0% AA), 0.05% AA, 0.5% AA, 1% AA and 1% CAW.

|  | Control (0) | 0.05% AA | 0.5% AA | 1% AA | 1% CAW | F, p |
| --- | --- | --- | --- | --- | --- | --- |
| Total distance traveled (m) | 11.55 +/- 2.38 | 15.53 +/- 1.52 | 15.60 +/- 1.85 | 16.36 +/- 1.92 | 15.08 +/- 1.32 | 1.11, 0.37 |
| Time immobile (log s) | 75.15 +/- 12.70 | 61.93 +/- 11.02 | 59.39 +/- 14.52 | 38.84 +/- 11.86 | 66.51 +/- 21.26 | 2.27, 0.78 |
| CFR time freezing (s) | 210.6 +/- 31.77 | 222.2 +/- 41.86 | 241.1 +/- 25.43 | 295.5 +/- 20.14 | 289.7 +/- 19.11 | 1.92, 0.12 |
| Synaptophysin (log fold induction) | 1.00 +/- 0.26 | 1.05 +/- 0.19 | 1.01 +/- 0.17 | 1.67 +/- 0.37 | 1.44 +/- 0.22 | 0.34, 0.85 |
| PSD95 (log – fold induction) | 1.00 +/- 0.15 | 0.93 +/- 0.14 | 1.11 +/- 0.18 | 1.66 +/- 0.42 | 1.58 +/- 0.28 | 0.34, 0.85 |
| Basal (pmol O2/min) | 127.57 +/- 3.66 | 131.13 +/- 4.26 | 142.78 +/- 4.56 | 153.31 +/- 8.43 | 143.74 +/- 4.94 | **3.65, 0.01** |
| Maximal (log pmol O2/min) | 198.07 +/- 12.53 | 195.93 +/- 8.92 | 215.09 +/- 8.55 | 235.94 +/- 11.99 | 232.59 +/- 26.22 | **2.72, 0.04** |
| ATP-linked (pmol O2/min) | 36.05 +/- 2.28 | 351.13 +/- 1.88 | 35.77 +/- 2.20 | 42.61 +/- 4.10 | 35.51 +/- 1.92 | 1.42, 0.24 |
| Spare Capacity (log pmol O2/min) | 70.50 +/- 13.17 | 64.80 +/- 5.98 | 72.31 +/- 8.21 | 82.63 +/- 7.68 | 88.84 +/- 16.04 | 0.91, 0.47 |
| Mt-ND1 (log fold induction) | 1.00 +/- 0.23 | 1.16 +/- 0.30 | 1.46 +/- 0.20 | 2.53 +/- 0.69 | 1.65 +/- 0.41 | 1.14, 0.25 |
| Mt-CYB (log fold induction) | 1.00 +/- 0.14 | 1.13 +/- 0.14 | 1.82 +/- 0.31 | 2.22 +/- 0.44 | 1.94 +/- 0.24 | 2.08, 0.10 |
| Mt-CO1 (log fold induction) | 1.00 +/- 0.42 | 1.17 +/- 0.44 | 1.30 +/- 0.38 | 1.85 +/- 0.36 | 1.61 +/- 0.52 | 1.93, 0.13 |
| Mt-ATP6 (log fold induction) | 1.00 +/- 0.24 | 1.16 +/- 0.37 | 1.88 +/- 0.25 | 2.77 +/- 0.73 | 1.96 +/- 0.27 | **4.43, 0.01** |
| NRF2 (fold induction) | 1.00 +/- 0.16 | 1.36 +/- 0.40 | 1.58 +/- 0.49 | 2.41 +/- 0.45 | 1.75 +/- 0.30 | **2.8, 0.04** |
| HMOX1 (fold induction) | 1.00 +/- 0.18 | 1.54 +/- 0.34 | 1.84 +/- 0.32 | 2.35 +/- 0.28 | 1.78 +/- 0.24 | **2.89, 0.05** |
| GCLC (fold induction) | 1.00 +/- 0.15 | 1.40 +/- 0.45 | 1.63 +/- 0.15 | 2.49 +/- 0.47 | 1.95 +/- 0.34 | **2.8, 0.04** |

### AA treatment increases contextual-associative memory in a dose-dependent manner

Contextual associative memory was evaluated with the CFR test. In the CFR, the animal can freely explore a chamber after which it is exposed to a mild foot shock. The following day the animal is reintroduced to that same chamber and the amount of time that the animal spends frozen is recorded over two five-minute periods. If the mouse remembers the association between the painful stimuli and the chamber, the amount of time frozen will be higher. Because we did not observe any differences in freezing between groups in either 5-minute recording periods the data presented is for the entire 10-minute window. There were no significant group differences between AA treated mice, CAW treated mice and control animals (Table 1); however, we did observe a significant linear relationship between increasing AA concentration and reduced time freezing in CFR performance (Figure 4A).

We measured the expression of the synaptic genes synaptophysin and PSD95 in cortical synaptosomes isolated from the brains of treated animals. There were no differences between any of the treatment groups (Table 1) nor was there a significant linear relationship between expression of these genes and concentration of AA (Figure 4B).

### AA alters cortical mitochondrial bioenergetics

The bioenergetics profile of cortical synaptosomes isolated from treated animals was analyzed using the SeahorseXF platform. There was a significant linear relationship between AA concentration and both basal and maximal mitochondrial respiration (Figure 5). Additionally, we found significant group differences in these endpoints (Table 1). Post-hoc analyses showed 1% AA significantly increased basal respiration relative to control treated mice. A similar but non-significant increase in maximal respiration between 1% AA treated mice and control mice was observed (p=0.10). CAW treatment did not result in significantly increased basal or maximal respiration.

**Figure 5.**
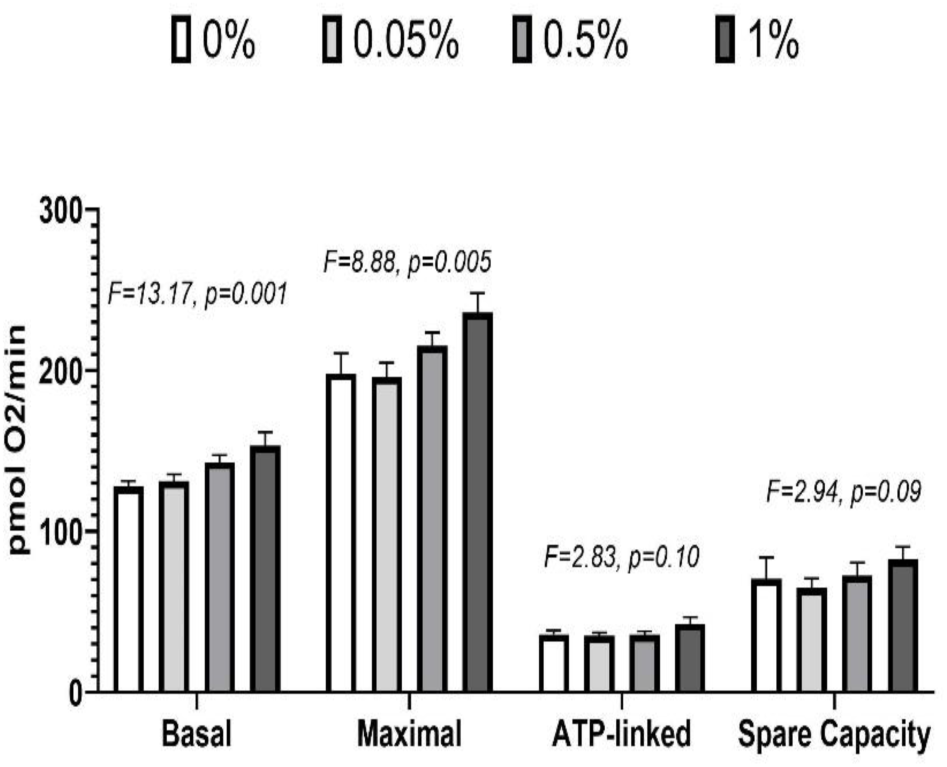
AA dose dependently improves basal and maximal mitochondrial respiration in cortical synaptosomes. Both basal and maximal respiration showed significant linear trends with AA concentration. No significant linear associations were found for ATP-linked respiration or spare capacity

AA treatment did not affect either ATP-linked respiration or spare capacity. There were no significant linear trends (Figure 5) or significant group differences (Table 1) for ATP production or spare capacity, a metric that reflects the extra energy available for demand increases that occur when cells are subjected to stress.

### AA dose dependently increases the expression of mitochondrial genes in the cortex of treated mice

We assessed the expression of the mitochondrial genes Mt-ND1, Mt-CYB, Mt-CO1 and Mt-ATP6 (which encode electron transport chain (ETC) complexes I, III, IV and V respectively) in cortical synaptosomes isolated from the brains of treated animals.

All four ETC genes showed a statistically significant linear trend of increased expression with increasing AA concentrations (Figure 6A).However, only Mt-ATP6 yielded statistically significant group differences (Table 1). Post-hoc analyses showed the 1% AA group had significantly higher Mt-ATP6 expression than both the control group and the 0.05% AA group.

**Figure 6.**
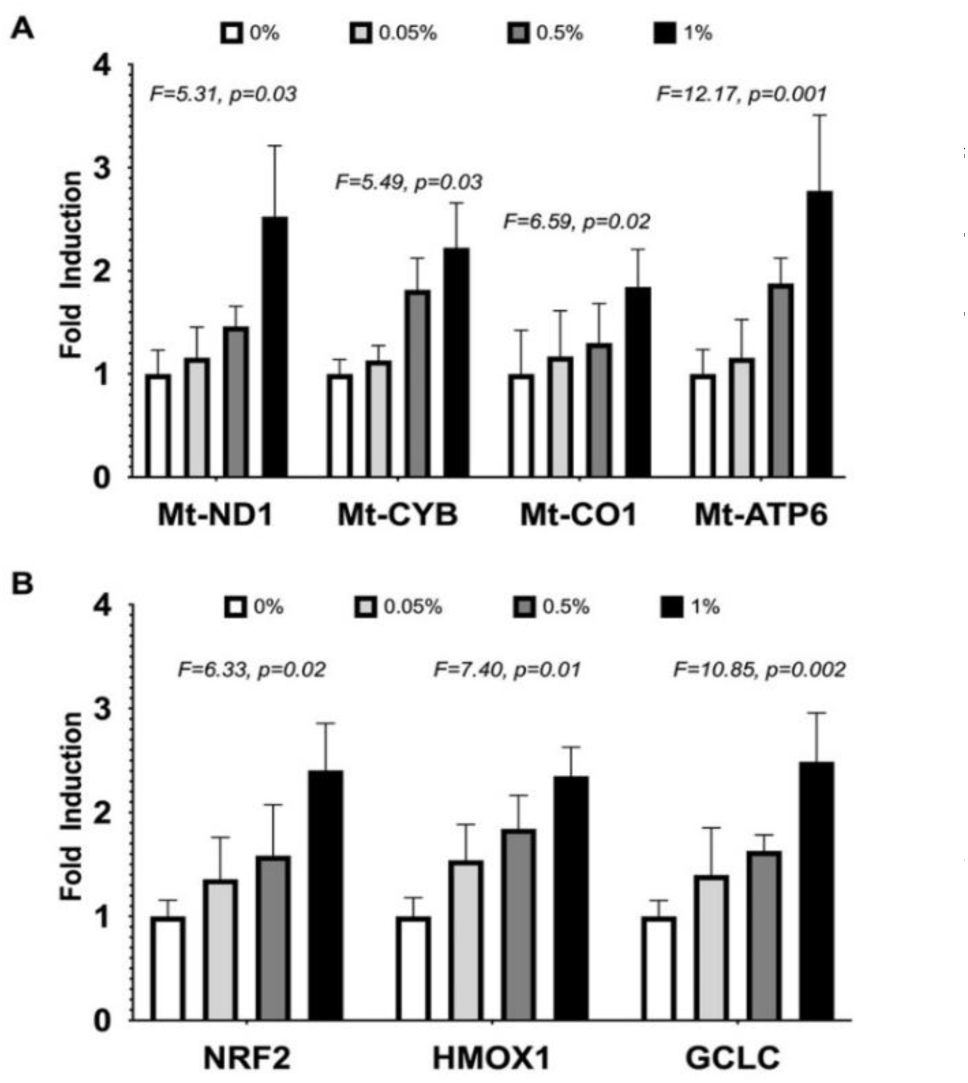
AA dose dependently increases the expression of mitochondrial and antioxidant genes. A) All four mitochondrial expressions (Mt-ND1, Mt-CYB, Mt-CO1 and Mt ATP6) in cortical synaptosomes showed statistically significant linear trends with increased AA concentration. B) All three antioxidant gene expressions (NRF2, HMOX1 and GCLC) in cortical synaptosomes showed statistically significant linear trends with increased AA concentration.

There was a similar, but non-significant trend towards an increase in Mt-ATP6 expression in the CAW-treated mice as well (p=0.14)

### AA dose increases the expression of antioxidant genes in the cortex of treated mice

We also evaluated the expression of antioxidant response genes in cortical synaptosomes isolated from treated animals. We measured expression of the antioxidant regulatory transcription factor NRF2 as well as its antioxidant target genes HMOX1 and GCLC in cortical synaptosomes. We observed a significant linear trend between increased expression of each of the three antioxidant response genes and increasing AA concentration (Figure 6B). Although statically significant models were observed for testing group differences (Table 1), the only statistically significant post-hoc pairwise comparison was in GCLC expression showing that mice treated with 1% AA had significantly higher expression than control animals. Again there was no difference in antioxidant gene expression between the control animals and CAW-treated animals.

### AA treatment improves associative memory in female 5xFAD mice

In the next phase of the study we treated male and female 5xFAD mice with 1% AA for 4 weeks and evaluated the same endpoints as in the dose response study. Model significance was tested using Kruskal-Wallis tests followed by Dunn’s test for post-hoc pairwise comparisons. 5XFAD female mice displayed a significant deficit in CFR performance compared to WT mice in both the first and second five-minute stage of the CFR test (χ²=9.89, p=0.02; χ²=9.21, p=0.03, respectively). AA treatment significantly improved CFR performance for the female 5xFAD mice in the first 5 minutes of the test (Figure 7A) and a similar trend was seen in with AA treatment in the second 5 minutes for the female 5xFAD mice as well (Figure 7B). No differences in freezing were observed in male mice regardless of genotype or treatment (Figure 7C (χ²=1.53, p=0.68) and 7D (χ²=0.46, p=0.93)).

**Figure 7.**
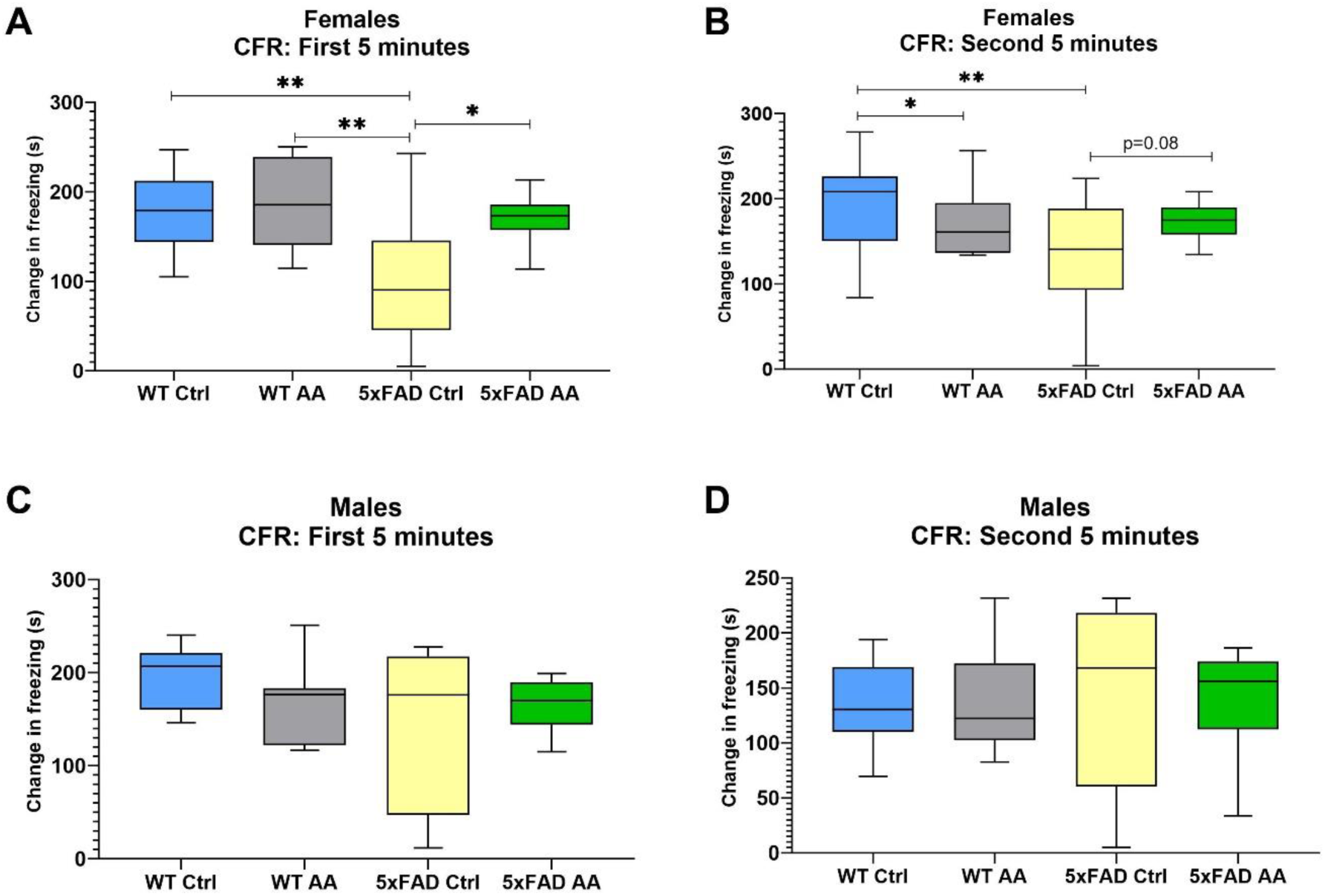
AA treatment changes time frozen in female 5xFAD mice. A) A significant improvement in CFR performance is seen in AA treated 5xFAD female mice over 5xFAD controls in first 5 minutes of test. B) Female 5xFAD AA treated mice see improvement in CFR score in second 5 minutes of test (p = 0.08). Female 5xFAD control mice have a significantly lower change in freezing compared to WT. C) Male 5xFAD and WT AA treated and control mice do not have significantly different changes in CFR performance in the first 5 minutes of the test. D) All 5xFAD and WT treatment groups do not have significantly different changes in CFR performance in the second 5 minutes of the test *p<0.05, **p<0.01

### AA treatment does not alter Aβ plaque burden

Brain tissue from 5xFAD mice was immunostained to quantify Aβ plaque pathology. We found that AA treatment did not alter plaque burden in either the hippocampus (Figure 8A (χ²=3.65, p=0.16) and B (χ²=0.88, p=0.35)) or cortex (Figure 8C (χ²=2.92, p=0.23) and D (χ²=0.10, p=0.75)) of either female or male 5xFAD mice.

**Figure 8.**
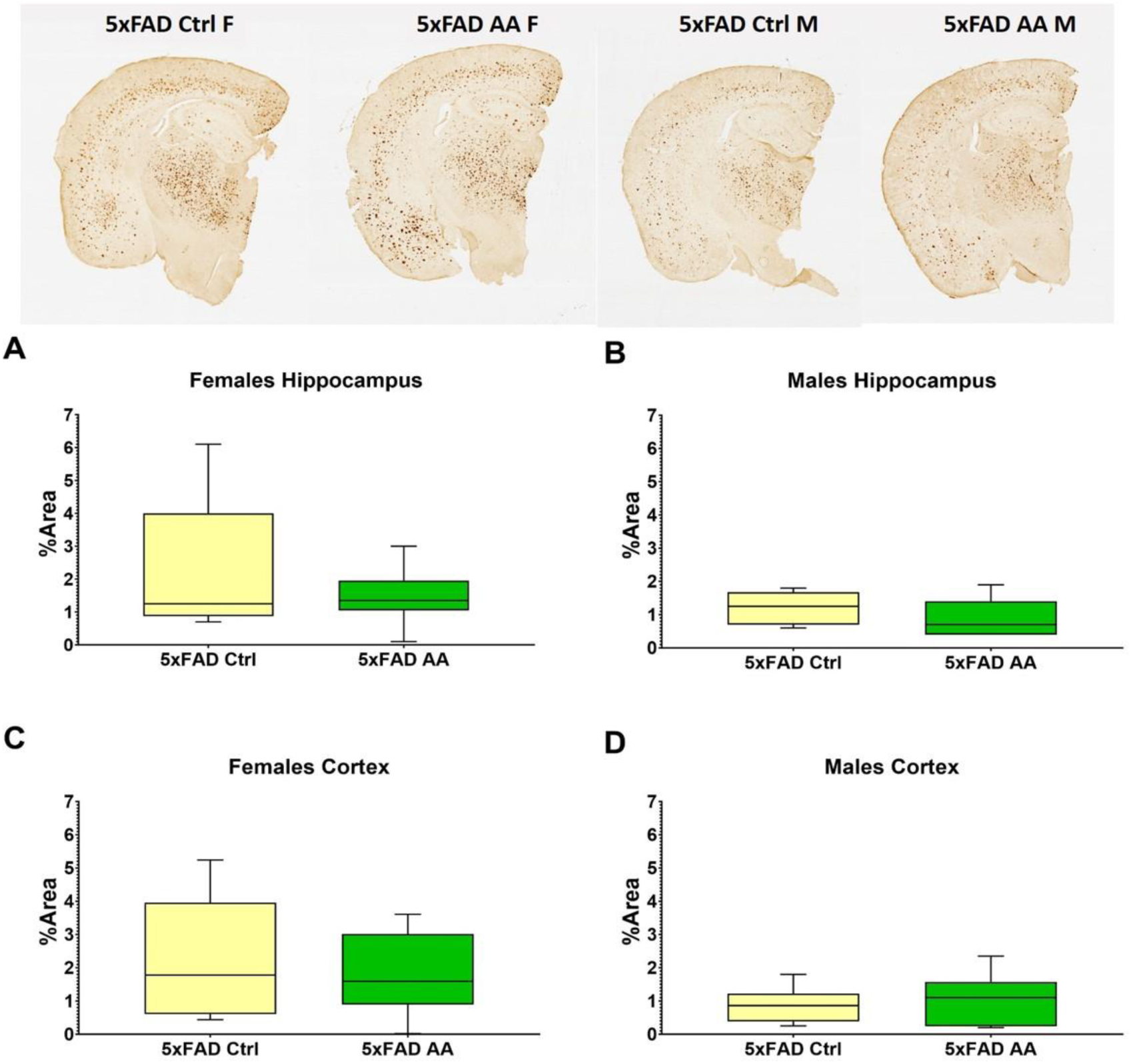
AA treatment does not alter Aβ plaque burden. Aβ plaque levels were not changed by AA treatment in either hippocampus (A,B) or cortex (C,D) of 5xFAD mice.

### AA treatment attenuates deficits in synaptic gene expression in female 5xFAD mice

We observed a reduction in the expression of synaptophysin in cortical synaptosomes isolated from 5xFAD female mice (Figure 9A). AA treatment significantly increased synaptophysin expression in female 5xFAD compared to 5xFAD controls (Figure 9A; χ²=8.41, p=0.04). Bordering statistical significance, a similar trend was observed with AA-treatment in male 5xFAD mice (Figure 9B; χ²=7.41, p=0.06). No change in PSD95 expression were observed in either male or female mice regardless of genotype or treatment (Figure 9C (χ²=2.52, p=0.47) and 9D (χ²=4.91, p=0.18)).

**Figure 9.**
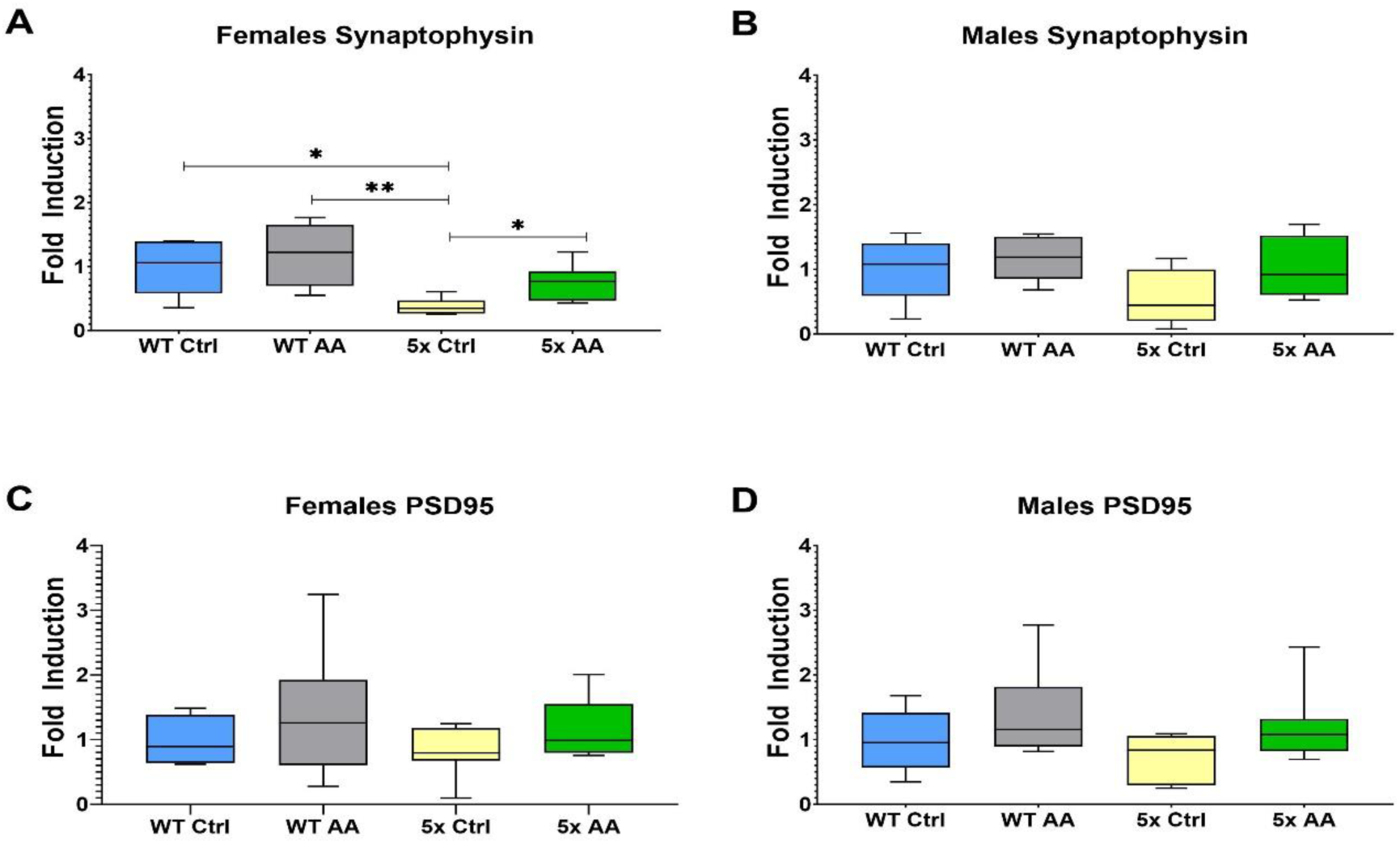
Cortical synaptophysin expression is increased in treated 5xFAD female mice. A) Synaptophysin expression levels in cortical synaptosomes are significantly decreased in female (A) and male (B) 5xFAD relative to WT controls. AA treatment significantly increased synaptophysin expression in female 5xFAD and had a similar but non-significant effect in male 5xFAD mice. C) No significant changes in PSD95 expression were observed in female (C) or male (D) mice regardless of genotype. *p<0.05, **p<0.01

### AA treatment improves mitochondrial bioenergetics in 5xFAD mice

A significant reduction in basal mitochondrial respiration was observed in cortical synaptosomes isolated from female 5xFAD mice relative to WT mice which was attenuated by AA treatment (Figure 10A; χ²=15.48, p=0.002). There was also a significant trend towards diminished maximal respiration evident in female 5xFAD mice that was likewise significantly increased in AA-treated 5xFAD female mice (Figure 10B; χ²=15.69, p=0.001). ATP-linked respiration and spare capacity were also assessed in cortical synaptosomes isolated from these mice. In female mice ATP-linked respiration was reduced in 5xFAD mice compared to wild type, and significantly increased with AA administration (Figure 10C; χ²=10.33, p=0.02). Increased spare capacity in AA-treated WT and 5xFAD female mice approached but did not reach significance (Figure 10D; χ²=7.57, p=0.06).

**Figure 10.**
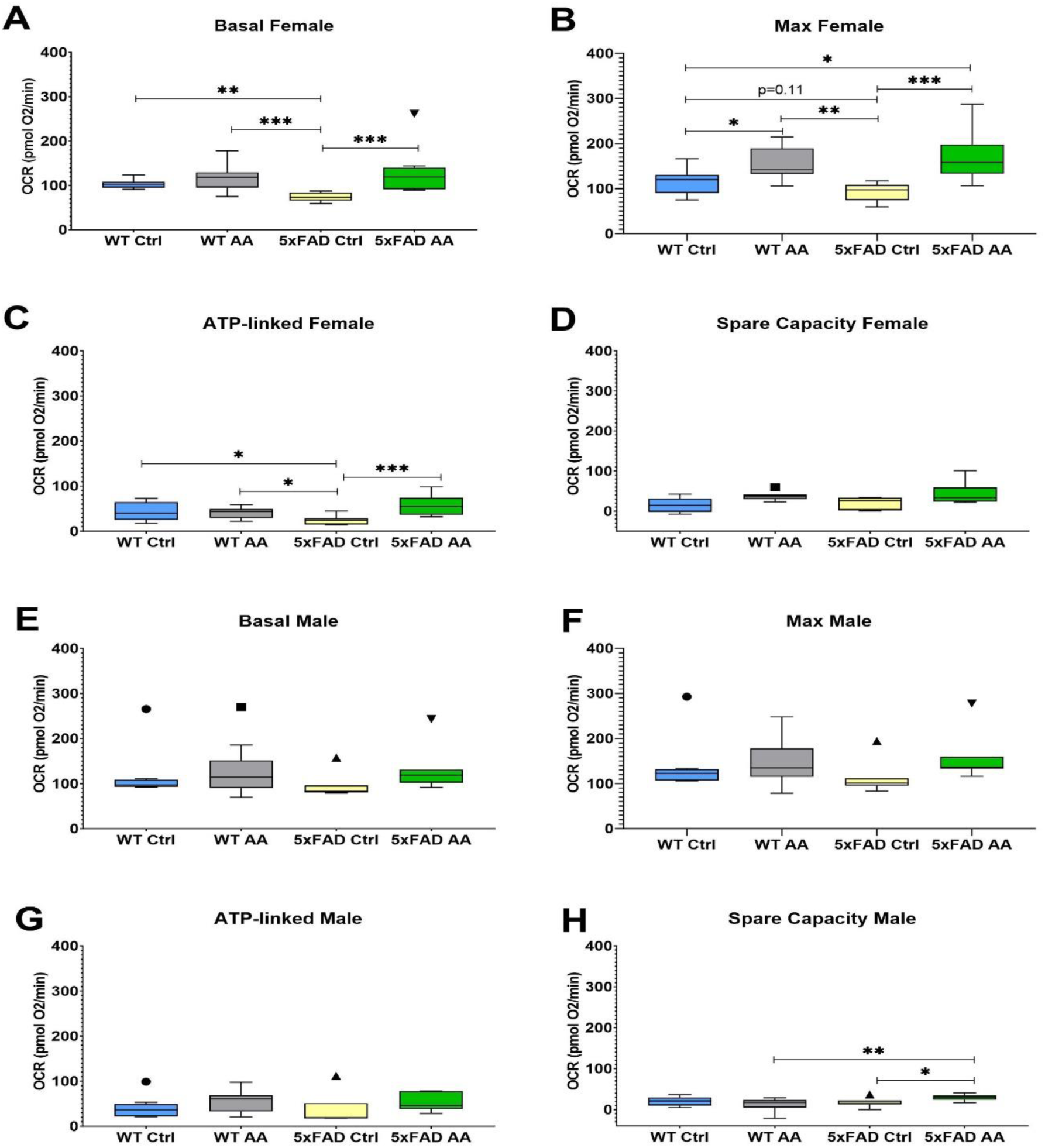
AA treatment improves cortical mitochondrial bioenergetics in 5xFAD mice. AA treatment attenuated deficits in basal (A), maximum (B) and ATP-linked (C) respiration in cortical synaptosomes from female 5xFAD mice. A similar but non-significant trend was also observed for spare capacity (D). In male mice deficits were not observed between 5xFAD and WT mice for any of the bioenergetic metrics (E-H) although an increase in maximal respiration (F) and spare capacity (H) was seen in AA treated 5xFAD male mice. *p<0.05, **p<0.01, ***p<0.001

The mitochondrial effects of AA were less pronounced in male mice. No significant changes in basal (Figure 10E; χ²=6.42, p=0.09) or ATP-linked (Figure 10G; χ²=5.64, p=0.13) respiration were observed in male mice regardless of genotype or treatment. There was, however, a statistically significant increase following AA treatment in spare capacity (Figure 10H; χ²=8.03, p=0.045) and a borderline significant increase in maximal respiration (Figure 10F; χ²=7.44, p=0.06) in 5xFAD male mice.

### AA increases expression of ETC genes in female 5xFAD mice

The expression of the ETC genes Mt-CYB, Mt-CO1 and Mt-ATP6 was reduced in cortical synaptosomes isolated from female 5xFAD mice as compared to WT mice (Figure 11B (χ²=9.54, p=0.02), 11C (χ²=8.96, p=0.03) and Figure 11D (χ²=9.20, p=0.03)). There was no change in the expression of Mt-ND1 in female 5xFAD mice relative to WT (Figure 11A; χ²=5.13, p=0.16). AA treatment robustly increased the expression of Mt-CYB, Mt-CO1 and Mt-ATP6 in female 5xFAD mice but did not affect the expression of any ETC genes in female WT mice.

**Figure 11.**
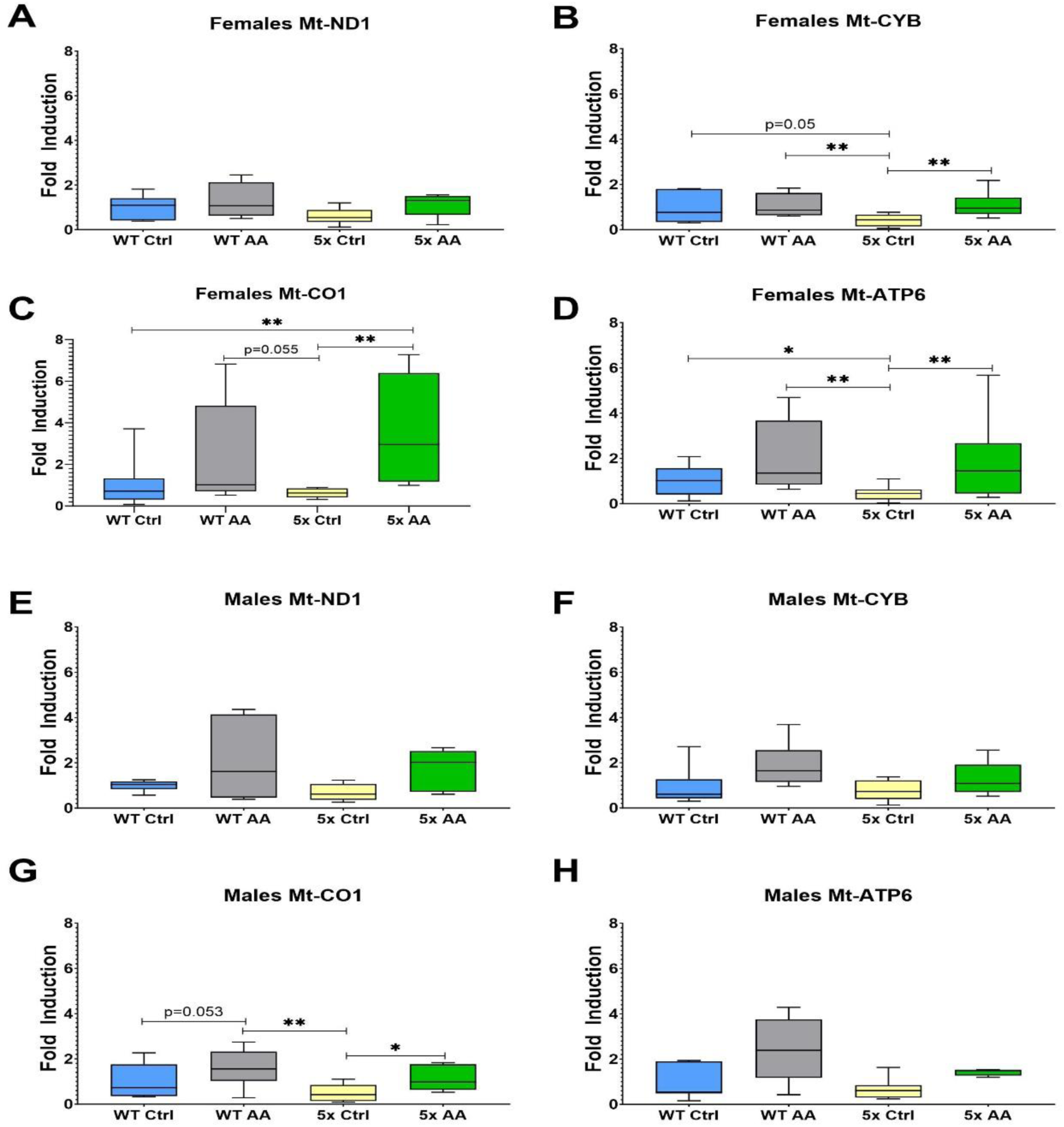
AA treatment induces cortical expression of ETC genes in female 5xFAD mice. ETC gene expression in cortical synaptosomes was quantified in female (A-D) and male (E-H) mice. AA treatment increased expression of Mt-CYB (B), Mt-CO1 (C) and Mt-ATP6 (D) in female 5xFAD mice. In male mice AA treatment only significantly increase expression of Mt-CO1 (G). *p<0.05, **p<0.01

In contrast, deficits in ETC gene expression were not observed in male 5xFAD mice relative to WT mice in Mt-ND1 (Figure 11E; χ²=5.82, p=0.12), Mt-CYB (Figure 11F; χ²=6.59, p=0.09), or Mt-ATP6 (Figure 11H; χ²=6.90, p=0.08). However, there was a statistically significant association in Mt-CO1 (Figure 11G; χ²=8.51, p=0.04). The only ETC gene affected by AA treatment in male 5xFAD mice was Mt-CO1.

### AA induces expression of antioxidant genes in female 5xFAD mice

The expression of the antioxidant regulatory transcription factor, NRF2, and its target antioxidant genes was increased with AA treatment in cortical synaptosomes from female 5xFAD mice as compared to control female 5xFAD mice (Figure 12A (χ²=7.84, p=0.05), Figure 12B (χ²=11.92, p=0.008), Figure 12C (χ²=10.02, p=0.02)). There was no effect of genotype or AA treatment on the expression of NRF2 and GCLC in male mice (Figure 12D (χ²=7.02, p=0.07), Figure 12E (χ²=2.5, p=0.47). Interestingly, there were differences in the expression of HMOX1 in male mice (Figure 12F; χ²=8.35, p=0.04), although they were related to genotype and not AA treatment.

**Figure 12.**
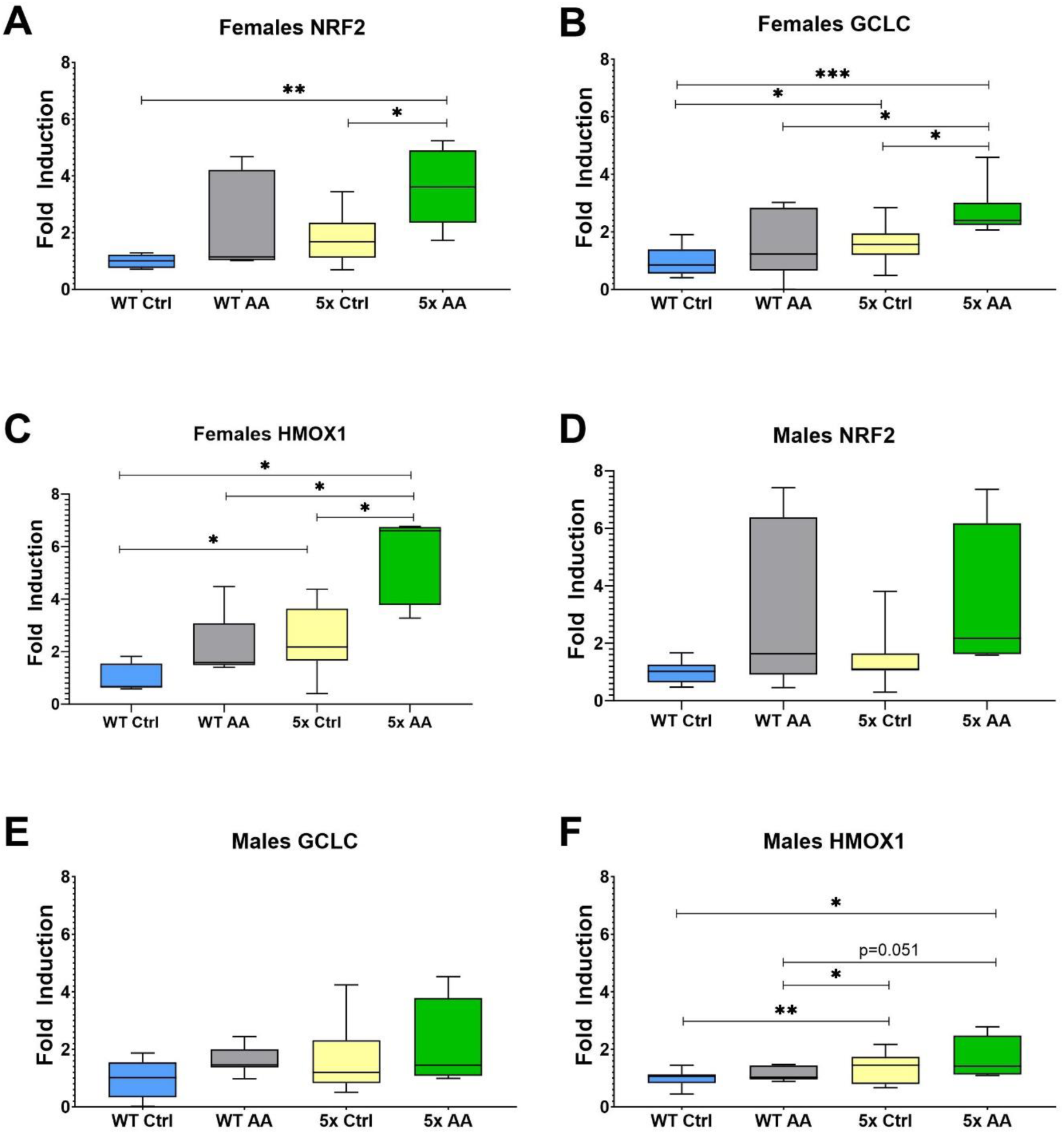
AA treatment induces cortical expression of ARE genes in female 5xFAD mice. AA treatment increased the expression of NRF2 (A) and its target genes GCLC (B) and HMOX1 (C) in female 5xFAD mice. HMOX1 expression was also significantly higher in female WT mice treated with AA than WT controls (C). In male mice there was no significant difference in NRF2 (D) and GCLC (E) between AA treated and control mice regardless of genotype. HMOX1 expression was significantly increased in AA treated WT mice relative to control WT mice (F). *p<0.05, **p<0.01

## Discussion

*Centella asiatica* has been used in traditional Ayurvedic medicine for over 3000 years and is widely recognized for its cognitive enhancing applications [47]. Both preclinical and clinical studies have shown therapeutic promise in improving cognitive function [16]. The herb extract is a complex chemical mixture containing several bioactive triterpenes and the precise compounds that are most responsible for the neuroprotective and cognitive enhancing effects of *Centella asiatica* have yet to be determined. In this study we have explored the effects of AA, the primary triterpene of *Centella asiatica,* in both healthy mice as well as in the 5xFAD model of Aβ accumulation. In the initial phase of this study, we found that AA dose dependently increased the cortical expression of antioxidant response genes and improved cortical mitochondrial bioenergetics in healthy mice. The effects of a high dose of AA (1% in the diet) administered to healthy mice elicited similar enhancements in both mitochondrial function and antioxidant gene expression as were evoked by a water extract of *Centella asiatica,* CAW. We went on to evaluate the effects of that high dose of AA in 5xFAD mice where in female mice we again observed increases in brain mitochondrial function and antioxidant response that were accompanied by an improvement in cognitive performance and an increase in synaptic gene expression, without significantly altering Aβ plaque burden.

We found that four weeks of AA treatment significantly increased the cortical gene expression of NRF2 regulated antioxidant enzymes in both healthy CF1 and Aβ-overexpressing 5xFAD mice. The NRF2 pathway is an important regulator in the antioxidant response. Oxidative stress is widespread in the AD brain [5] and upregulation of NRF2 has been shown to ameliorate oxidative damage [48]. The effects of AA on NRF2 observed in the present study are in line with previous findings. *In vitro*, AA treatment of HepG2 cells challenged with tert-butyl hydroperoxide reduced ROS accumulation and apoptosis via NRF2 upregulation [49]. Similar beneficial effects of NRF2 activation by AA have been observed *in vivo*. Our own lab has previously reported increases in antioxidant response gene expression in the brains of both aged healthy and 5xFAD mice treated with the AA-containing CAW extract [11, 12, 17, 21]. Other groups have likewise found that AA increases NRF2 expression and reduces markers of oxidative stress in rodent models of neurological injury including doxorubicin-induced toxicities, spinal cord injury and traumatic brain injury [50–52].

A reduction in the number of mitochondria is evident in the AD brain [9]. The fact that in the present study AA increased mitochondrial gene expression suggests that AA could be attenuating this loss of mitochondrial density. This may be due to an effect on mitochondrial biogenesis. While to our knowledge this possibility has not been evaluated *in vivo, in vitro*, treatment of a human neuroblastoma cell line with AA resulted in enhanced expression of peroxisome proliferator-activated receptor γ coactivator α (PGC-1α), an important regulator of mitochondrial biogenesis [53]. Future studies are needed to confirm an effect of AA on mitochondrial content *in vivo to* determine if the mechanism of action of AA includes mitochondrial biogenesis.

In this study we also found the AA treatment improved mitochondrial bioenergetics in the brains of treated mice. These results are similarly consistent with our group’s prior studies evaluating CAW which found increased expression of ETC genes in the hippocampus of healthy aged and Aβ overexpressing mice [11, 17]. There are other reports in the literature of mitoprotective effects of AA specifically as well. AA has been shown to protect mitochondrial membrane potential in rodent liver cells [54, 55] and in a mouse model of stroke AA treatment resulted in reduced cortical mitochondrial dysfunction [56].

Although to our knowledge this is the first time AA has been evaluated in a mouse model of Aβ accumulation, the cognitive enhancing effects of AA that we observed in 5xFAD mice have been well-documented in previous studies by our group and others. We have shown that CAW treatment increases synaptic gene expression and improves cognitive function in Aβ overexpressing mice [12, 17, 21, 22]. In a rat epilepsy model of cognitive impairment, pretreatment with AA increased levels of synaptophysin in the hippocampus and improved deficits in learning and memory [57]. Other structurally similar constituent triterpenes from *Centella asiatica* have been reported to similarly attenuate cognitive deficits. Asiaticoside, the metabolic precursor of AA, has been shown to improve memory in senescence accelerated mice and madecassoside, has been reported to improve memory deficits caused by both lipopolysaccharide and D-galactose [25, 30]. Moreover, the cognitive enhancing effects of other polyphenol compounds, including from coffee, grapes, and blueberries, have likewise been demonstrated in multiple mouse models of cognitive impairment including models of Aβ accumulation as well as in healthy and cognitively impaired human populations [58–63]. Interestingly, the effects of AA were consistently more pronounced in female 5xFAD mice than in males across virtually all of the endpoints measured. This variability in response between sexes is in line with our previous studies of CAW in 5xFAD models where we also observed a greater response to CAW in female animals [12, 21]. One possible explanation for these sex difference could be related to the greater plaque burden seen in 5xFAD female mice. The greater plaque burden in female 5xFAD mice is associated with more pronounced downstream effects such as cognitive and mitochondrial impairments which makes it easier to detect a treatment effect than in male 5xFAD mice where those deficits are more subtle.

Another potential explanation for the sex differences observed could be related to differential actual exposure of AA. All mice were exposed to the same concentration of AA in the diet but the fact that female mice are smaller than male mice suggests that they may in fact receive a higher exposure in terms of mg/kg bodyweight. Unfortunately, in this study mice were group housed and therefore individual diet consumption was not measured. Future work is needed with more rigorously controlled dosing to fully understand the potential sex differences in bioavailability and effects AA.

It is notable that in our study AA treatment improved cognitive, mitochondrial and antioxidant endpoints without altering plaque burden in the 5xFAD mice. Again, this finding is consistent with our previous CAW studies in Aβ overexpressing mice where we did not see a change in Aβ levels following treatment [21, 22]. This suggests that the improvements we saw following AA treatment are independent of plaque levels per se and that the intervention is instead targeting the downstream consequences of the plaques. If this is in fact the case, then AA could prove very promising for therapeutic development especially given the limited implementation and effects of Aβ targeting therapies across the AD community unsuccessful [64].

In conclusion, the results from this study suggest that oral AA administration can enhance mitochondrial function, induce antioxidant response, and improve cognitive function in Aβ overexpressing mice. Future studies are needed to optimize the timing of dosing and confirm in other AD models to fully explore its potential as an AD therapeutic agent. Moreover, since mitochondrial dysfunction and oxidative stress accompany cognitive impairment in other neurodegenerative diseases, the effects of AA in conditions beyond AD also warrant further investigation.

## Acknowledgments

This research was funded by a VA merit review grant to JQ.

